# Selection of optimal cell lines for high-content phenotypic screening

**DOI:** 10.1101/2023.01.11.523662

**Authors:** Louise Heinrich, Karl Kumbier, Li Li, Steven P. Altschuler, Lani F. Wu

## Abstract

High-content microscopy offers a scalable approach to screen against multiple targets in a single pass. Prior work has focused on methods to select “optimal” cellular readouts in microscopy screens. However, methods to select optimal cell line models have garnered much less attention. Here, we provide a roadmap for how to select the cell line or lines that are best suited to identify bioactive compounds and their mechanism of action (MOA). We test our approach on compounds targeting cancer-relevant pathways, ranking cell lines in two tasks: detecting compound activity (“phenoactivity”) and grouping compounds with similar MOA by similar phenotype (“phenosimilarity”). Evaluating six cell lines across 3214 well-annotated compounds, we show that optimal cell line selection depends on both the task of interest (e.g. detecting phenoactivity vs. inferring phenosimilarity) and distribution of MOAs within the compound library. Given a task of interest and set of compounds, we provide a systematic framework for choosing optimal cell line(s). Our framework can be used to reduce the number of cell lines required to identify hits within a compound library and help accelerate the pace of early drug discovery.

## Introduction

Large, diverse libraries of novel small molecules serve as critical components in the early drug discovery screening pipeline. ^1^ The cellular activities of the compounds in these libraries are typically unknown. High-content microscopy is a scalable approach for characterizing the effect of small molecules on cells. ^2–5^ In this setting, cellular responses to compounds are represented as feature vectors, whose entries measure observable changes, such as in cell morphology, biomarker intensity and localization. A single-pass phenotypic profiling screen can provide annotation to compound libraries by identifying subsets of library compounds that show “phenoactivity” (i.e., induce cellular responses distinct from control conditions) and by inferring MOA through “phenosimilarity” (i.e., by comparing cellular responses of compounds annotated with the same MOA). ^6–9^

Phenotypic profiling screens depend critically on the selection of cellular readouts and screened cell lines. For the purposes of annotating large compound libraries, previous work has focused on selection of “optimal”^10^ as well as general^11^ combinations of cellular readouts for a given cell line. However, approaches for selecting optimal cell lines are less explored. ^12–14^ It is reasonable to expect that different cell lines may have different sensitivities to detect different MOAs. For annotating diverse compound libraries in an unbiased- or target-agnostic fashion, it is poorly understood how to select the best performing cell line and to what extent using multiple cell lines would improve coverage.

To explore these questions, we first generated a high-content microscopy dataset of six cell lines responding to a diverse set of 3214 small molecules with annotated MOA. We developed computational methods to rank cell lines and combinations of cell lines according to their ability to infer compound activity and MOA. Lastly, we applied a classic optimization framework to assess which cell lines or cell line combinations would be best for annotating uncharacterized compounds.

## Results

### Experimental design

Our dataset was generated using six cell lines. These included five cancer cell lines (A549, OVCAR4, DU145, 786-O, HEPG2) from the NCI60 set of tumor cell lines; these cell lines span a range of tissue types (including epithelial, endothelial, neuronal, secretory (ductal), monoblast and erythroid origins), cellular morphologies, and are amenable to imaging assays. We additionally included a non-cancer patient-derived fibroblast cell line (“FB”). Additional details in Supporting Information.

We collected a “reference” library of 3214 compounds annotated with mechanism of action (MOA). This library includes FDA-approved drugs, in-clinical-trial drugs, and established bioactive tool compounds; these compounds cover 664 MOAs (Supporting Information Table 1) in both cancer-related pathways as well as the broader druggable target space. Due to the number of cell lines and drugs, we limited data collection to 48h, and a single dose (either 5uM or 10uM, depending on the library used). We selected a 48h time point, taking into account that large screens of diverse compound sets are likely to include compounds that act over different time frames. We previously observed that this time point provided high classification accuracy on reference drug classes and overall diversity of phenotypic responses.^10^

To label cells, we made use of the Cell Painting assay^11^ and sampled nine fields of view at 20X magnification for each well by microscopy. For each treatment, we performed cell segmentation and feature extraction to generate 77 quantitative features (Supporting Information Table 2) describing cellular morphology and the distribution (i.e. intensity, texture, objects) of the six intracellular stains that comprise the Cell Painting marker set. We generated phenotypic profiles that summarized the population-level shift in each feature from a negative control condition (DMSO) using signed KS statistics^7^ (Supporting Information). Thus, the cellular response to each compound perturbation was summarized as a 77-dimension profile in phenotypic space.

### Data analysis

Using these data, we sought to identify “optimal” cell lines or combinations of cell lines using two measures of annotation performance on the reference compound library: (1) *phenoactivity* : the degree to which compound and DMSO profiles differ; and (2) *phenosimilarity* : the degree to which the compounds with the same MOA share similar phenotypic profiles (Supporting Information).

Our approach focused on three key challenges. First, quantification of phenoactivity and phenosimilarity depend on analytical parameters, which can be challenging to tune across multiple cell lines. We aimed to limit the number of such parameters and ensure that results were robust to these parameter choices. Second, cellular phenotypic changes from DMSO control can range from subtle to severe across different cell lines. We aimed to ensure that our measurements of phenoactivity and phenosimilarity were sensitive across this spectrum and could report on responses that were similar to, but distinct from control. Third, compounds annotated to share a common MOA may induce phenotypically dissimilar responses (e.g., due to different on- or off-target activities, potency, coarse annotation, and so on). We aimed to define “phenotypic tightness” of an MOA in a manner that was robust to some degree of compound heterogeneity. Based on these requirements, we chose to define phenoactivity and phenosimilarity at the MOA level rather than at a compound-by-compound level. In brief, for each cell and MOA, we computed: (1) a phenoactivity score by comparing the distributions of distances of the MOA and DMSO point clouds to the centroid of the DMSO point cloud; and (2) a phenosimilarity score by comparing the tightness of the MOA point cloud relative to the nearest neighbor point clouds of each MOA compound.

### Phenoactivity

We first assessed the ability of individual cell lines to detect phenoactivity of compounds from specific MOA classes (Supporting Information Table 3). In all cell lines, positive control compounds were phenotypically distinct from DMSO controls (Fig. S4). We visualized how compound profiles were distributed in phenotypic space for different cell lines using UMAP.^15^ In some cases, point clouds for collections of compounds annotated with the same MOA and DMSO replicates were similarly separated across cell lines, while in others dramatic cell line differences were apparent. For example, in the case of the HDAC inhibitors, both HEPG2 and OVCAR4 cell lines showed similar degrees of separation between HDACs and DMSO, with both cell lines detecting phenoactivity for a similar subset of HDAC compounds (e.g., 28/35 and 29/35 of HDAC compounds fell outside the DMSO point cloud—were more than one IQR above the median DMSO to DMSO centroid distance—in OVCAR4 and HEPG2 respectively; Fig. 2a, top). In contrast, in the case of glucocorticoid receptor agonists (GRA), for OVCAR4 all GRA compounds were outside the DMSO cloud while a minority were for HEPG2 (29/29 vs 11/29 for OVCAR4 vs HEPG2, resp.; Fig. 2a, bottom). These observations were consistent with our phenoactivity scores based on conversion of these point clouds into distance distributions (Fig. 2b). We summarized phenoactivity scores across all MOA-cell line pairs to provide an overview of differential activity across the entire compound library (Fig. 2c). While OVCAR4 was overall the most sensitive for detecting phenoactivity, another cell line performed better in 88/148 MOAs containing at least 5 compounds.

Interestingly, most MOAs showed low phenoactivity in HEPG2. To assess phenotypic features that most differentiated HEPG2 from other cell lines, we trained an iterative random forest^16^ to classify HEPG2 versus other cell lines treated with DMSO control (Supporting Information; Fig. S5). The classifier achieved near perfect accuracy on a hold-out test set of well replicates (area under the receiver operating characteristic curve 0.999), suggesting that HEPG2 cells were systematically distinct from the remaining cell lines (Fig. S5b). To identify features that drove this separation, we evaluated mean decrease in impurity (MDI) feature importance and found that the classifier was heavily influenced by cell nearest neighbor distance (Fig. S5c). This finding was consistent with qualitative examination of cell images (Fig. S5a), which highlighted HEPG2 cell’s tendency to clump closely together.

We hypothesize that the growth of HEPG2 in highly compact colonies underlies the poor performance of this cell line in producing phenotypic profiles that are able to distinguish compound-induced phenotypes from control. Several of the markers used in our study target cellular organelles (mitochondria, actin) that would be difficult to distinguish alterations in, in a compact cell. In addition, overall morphology (geometry) features of the cell are an important driver of phenotypic variation, and HEPG2 is less variable in its geometry due to its colony growth pattern. This example of a poorly performing cell line highlights the importance of considering cell morphology when selecting a cell line of interest for phenotypic screening—and exemplifies the use of our framework to select-out cell lines with these properties.

### Phenoactivity optimization

How much improvement in phenoactivity detection is provided by inclusion of additional cell lines? We compared the abilities of individual and pairs of cell lines to detect phenoactivity across all MOA categories (Fig. 2d). For a pair of cell lines, we defined the phenoactivity of an MOA as the maximum phenoactivity over each individual line—effectively asking whether phenoactivity is detected in either cell line (Supporting Information). The single best performing cell line is OVCAR4. By construction, pairs of cell lines including OVCAR4 outperform OVCAR4 on its own. However, the improvements relative to OVCAR4 were marginal (phenoactivity score 0.576 OVCAR4 vs. phenoactivity score 0.611 OVCAR4, 786-0; *∼* 6.1% increase).

### Phenosimilarity

We next assessed whether compounds with the same provided MOA induce similar cellular phenotypes, which we quantified through phenosimilarity scores (Supporting Information Table 4). As previously, we assessed the ability of individual cell lines to detect phenosimilarity of specific MOA classes (Fig. 3a-b). Here, we compared the distribution of distances among compounds within an MOA (colored distributions) to the distribution of those compounds’ nearest neighbors (gray distribution). In the case when an MOA is tightly clustered, the nearest neighbors of a compound will be another compound in the same MOA and the two distributions will closely overlap; conversely, if the point cloud of an MOA is broadly distributed and contains profiles of compounds in other MOAs, these distribution will look dissimilar. We note that the ability to detect phenosimilarity in a given cell line is dependent on the specific set of experimental parameters considered in our analysis (e.g. marker set, treatment time, treatment dose). Cellular heterogeneity may be driven in part by the fact that phenoactivity and phenosimilarity for a particular MOA / cell line combination cannot be detected with the parameters used in our analysis.

MOAs with high phenosimilarity in a given cell line were tightly clustered and fell further from the DMSO point cloud (Fig. 3a). MOAs with low phenosimilarity across all cell lines were evenly distributed across the DMSO point cloud (Fig. S6)—i.e. they could not be distinguished from DMSO controls. As a specific example, we compared MOA point clouds and phenosimilarity scores between the A549 and FB cell lines (Fig. 3a-b). While HSP inhibitors were tightly clustered in both cell lines (phenosimilarity score 0.828 and 0.747 in A549 and FB respectively). SYK inhibitors, glycogen synthase kinase (GSK) inhibitors, and tubulin polymerzation (TP) inhibitors showed varying degrees of differential clustering behavior between the two cell lines (SYK inhibitor phenosimilarity score 0.061 and 0.317 in A549 and FB respectively, GSK phenosimilarity score 0.386 and 0.102 in A549 and FB respectively, TP phenosimilarity score 0.538 and 0.428 in A549 and FB respectively).

TP inhibitors that were distinct from DMSO clustered more closely in phenotypic space for A549 compared to FB, reflecting its slightly higher phenosimilarity score. Phenotypically, these compounds induced changes in morphology that can be interpreted as the result of cell cycle defects, observed in response to microtubule inhibition at longer timepoints (e.g. 48h). This is a well-described phenotype resulting from microtubule inhibition in the literature.^17–19^ Specifically, upon treatment, we observe decreases in nuclear roundness and nuclear area, and corresponding increases in cell roundness, in addition to changes in the texture of signal intensity of the Hoechst (DNA) channel (Fig. S7). Differential phenosimilarity of GSK3 inhibitors between A549 and FB can be explained by the distinct roles of GSK3 in these 2 cell lines. Non-small cell lung cancer cell lines, such as A549, exhibit increased GSK3 kinase activity, which supports tumor cell proliferation and contributes to the poor prognosis.^20,21^

Thus, GSK3 inhibition is expected to produce phenotypes in A549. Indeed, 8 / 11 GSK3 inhibitors showed phenoactivity in A549, and clustered closely together in phenotypic space (only SB216763, Tideglusib, and SB415286 clustered near DMSO). In contrast to A549, increased GSK3 kinase activity has not been reported in the non-cancer FB cell line and GSK3 inhibitors showed low rates of phenoactivity and phenosimilarity.

Moderate phenosimilarity scores for mTOR inhibitors (0.511 A549, 0.586 FB) highlight the fact that while these compounds cluster near one another in phenotypic space, they closely neighbor other compounds with different MOA annotations. In the case of A549, a subset of closely grouped mTOR compounds is reflected in the bi-modality of the distance distribution (Fig. 3a-b). The two subclusters of mTOR inhibitors contain compounds with different inhibition mechanisms: one cluster is formed solely from mTOR kinase domain inhibitors that inhibit the activities of both mTORC1 and mTORC2, while the other cluster is formed mainly from allosteric inhibitors (rapamycin-analogues) that inhibit mTORC1 more selectively (Fig. 3e). Qualitative examination of phenotypes revealed that the mTOR kinase domain inhibitor group induced a “clumping” phenotype (Fig. 3f, INK 128 and Torin This phenotype is characterized by features capturing distorted morphology (e.g. cell compactness, radial axis length, contact with neighbors, etc.; Fig. 3e). In contrast, the allosteric inhibitor group showed less severe differences in morphology but induced changes in cellular objects (Fig. 3e).

### Phenosimilarity optimization

Which cell line is the best “generalist” for grouping compounds based on these provided MOAs? We summarized phenosimilarity scores across all MOA-cell line pairs (Fig. 3c). In this dataset, A549 was the most sensitive for grouping compounds with the same MOA, and thus was the best generalist. Nevertheless, there were 108/149 MOAs in which another cell line performed better. For screens focused on these specific MOAs, other cell lines may perform better as “specialists”. How much improvement in MOA phenosimilarity is provided by inclusion of additional cell lines? We computed the MOA phenosimilarity across a pair of cell lines by taking the maximum score across individual lines—effectively asking whether an MOA is tightly clustered with respect to either cell line (Supporting Information). Here, the best performing cell line based on phenosimilarity is A549. Including an additional cell line with A549 offered a *∼* 28% improvement in average phenosimilarity score (0.148 for A549 vs. 0.189 for A549, FB), driven by addition of several MOAs that were poorly clustered in A549 (e.g., FB improved identification of SYK, HMGCR, and proteasome inhibitors; Fig 3d).

## Discussion

A fundamental choice when setting up a large-scale high-content phenotypic screen is to determine which cell line or cell lines are best suited to identify bioactive compounds and predict their MOAs. Here, we provide a road map for how to objectively answer this question. We make use of an annotated reference compound library with hundreds of MOA classes to calibrate the performance of each potential screening cell line as well as combinations thereof. Phenoactivity scores can be used to select cell lines that best identify bioactive compounds. Phenosimilarity scores can be used to select cell lines that best identify compounds within specific MOA(s).

In our case study, we found that the best performing cell lines were different for either identifying bioactive compounds or categorizing compounds by MOA classes, and that the utility of using additional cell lines was not the same for each of these tasks. A cell line that is optimal to detect phenoactivity may have poor sensitivity for the same MOA class (and vice versa) (Fig. 3e). This is because the ability to infer MOA from clusters in phenotypic space requires that compounds within the target MOA are near one another yet far from compounds with other MOAs. Phenosimilarity scores capture these two properties. In practice, large compound libraries could be annotated by evaluating how phenotypically similar unknown compounds are to known reference classes. Phenosimilarity scores assess the degree to which reference classes are phenotypically consistent and thus provide a measure of confidence for the degree to which new, phenotypically similar compounds belong to a given MOA.

There are a number of ways in which our approach could be improved or extended. From a platform perspective, we made a number of technical choices, including compound dose, treatment time, cell features, construction of phenotypic profile, distance metric, optimization framework, and so on. Each of these choices can be further examined for improved performance. From a more general perspective of using high-content imaging to detect and predict MOA of uncharacterized compounds, three are three major inputs to this process: cell line, biomarker and annotated reference compound sets. Here, we optimized over cell lines while holding the biomarkers (cell painting) fixed, while in past work we optimized over biomarkers while holding a cell line fixed.^10^ In both cases we assumed that a reference compound library was provided with MOA annotation classes. A future task is to investigate how phenosimilarity scores can be used to refine or coarsen provided MOA annotation classes.

For instance, by replacing the DMSO reference distribution with some other MOA class in our phenoactivity analysis, one could evaluate the similarity between two MOA classes and provide guidance for when two MOA classes should be merged. Such analyses offer a new path to annotating compound libraries based on phenotypic consistency. A larger task, for future work, is an experimental and computational platform designed to optimize over all three input choices for a given screening goal.

High-content microscopy screens offer the promise to annotate large libraries of uncharacterized compounds. However, the ability of different cell models to classify compounds varies according to their underlying biology. Here, we show an objective framework for selecting one, or a small number of cell lines that are “optimal” for predicting bioactivities across specified MOA classes. It is our hope that this framework will help increase the scale, sensitivity and accuracy of phenotypic profiling used in early drug discovery.

### Data and code availability

Data and code to reproduce figures and analyses are available on [Zenodo](https://doi.org/10.5281/zenodo.7352487).

## Acknowledgement

We are grateful to members of the Altschuler and Wu labs for constructive feedback. We gratefully acknowledge NCI-NIH RO1 CA184984 (L.F.W. and S.J.A.).

## Supporting Information

## Materials and methods

### Compound library

All screening plates were treated with 28 wells of vehicle (DMSO), and 4 wells containing positive controls (Bortezomib 5uM, Gemcitabine 5uM). Compound libraries were sourced from Selleck Chemicals LLC: FDA-approved & Passed Phase I Drug Library (L3800) (10uM), Kinase Inhibitor Library (L1200)(10uM), Apoptosis Compound Library (L3300)(10uM), Epigenetics Compound Library (L1900) (10uM),and Bioactive Library (L1700)(5uM). In addition to these compound libraries, in each screening batch we also added two replicate plates containing a “Reference Set” of 35 compounds, each present at 5 5-fold dilutions as described in.^1^

### Compound MOA annotation

Screened compounds with the same chemical structure (SMILES and InChIKey) are combined under the same common name (Compound ID). MOA annotations are curated from the Drug Repurposing Hub database and mapped to each compound based on their chemical structures. For a full list of compounds screened and annotations assigned (Supporting Information Table 1).

### Cell culture

We cultured a subset of the NCI60 database, including A549, 786-O, OVCAR4, DU145 and HEPG2. In addition, we added a healthy fibroblast line “FB”. A549, 786-O, OVCAR4, DU145 were cultured in RPMI1640 (Thermo Fisher Scientific #21875-034), supplemented with 10% heat-inactivated FBS (Gemini #CAT) and 1X antibiotic-antimycotic (Thermo Fisher Scientific #15240062). HEPG2 were cultured in DMEM (Thermo Fisher Scientific A4192001) supplemented with 10% heat-inactivated FBS (Gemini #100-500) and 1X antianti (Thermo Fisher Scientific #15240062). FB were cultured in DMEM/F-12 (Thermo Fisher Scientific, #11320-033), 20% FBS, 1% GlutaMAX (2 mM), 1% Pen-Strep (Gibco #15140122). Cells were plated in 384-well plates (Perkin-Elmer Cell Carrier Ultra #6057300) in 75uL media. 1000 cells/well for A549, OVCAR4, DU145, 786-0 and FB. HEPG2 were plated at 2500 cells/well.

### Compound treatments

Compound treatments were added using an Echo 650 (Beckman Coulter), to a final DMSO concentration of 0.1% (75nL compound or vehicle in 75uL media). Compounds were added at either 10uM or 5uM (library dependent). Cells were incubated with compound for 48h.

### Microscopy

Plates were imaged in confocal mode on the Operetta CLS high-content imaging system (Perkin Elmer) using a 20X water immersion lens (NA1.0), effective resolution (0.66um). 9 fields of view were captured per well using a 4.7MP 16-bit sCMOS sensor (6.5um pixel size).

### Feature extraction

Harmony (version 4.9, Perkin Elmer) software was used to segment individual cells and extract features. First, images were flatfield-corrected and background subtracted. Next, individual nuclei were segmented using the Hoechst channel, and cytoplasm using the Alexa 568 channel (WGA, Phalloidin). Cells touching image borders were filtered out. We next calculated groups of features encompassing morphology, intensity and texture for each channel, totalling 76 features. For a full list of features, see Supporting Information Table 2.

### Phenotypic profiles

Phenotypic profiles transform the measured features of single cells within a treated well into a population-level measure of the deviation from the negative controls contained in the same plate, using a non-parametric signed Kolmogorov-Smirnov (KS) statistic.^2^

### Plate position normalization

Preliminary data analysis showed positional effects for a subset of plates in our dataset. That is, certain rows/columns of the plate showed systematically higher/lower feature levels among DMSO controls. These effects were also observable in PCA projections of DMSO controls (Supporting Information Fig. S3). To address these effects, we regressed phenotypic profiles against row and column IDs within each plate and subtracted out predicted values, effectively removing the portion of a phenotypic profile that could be explained by plate position (Supporting Information Fig. S1-S3). Following this regression, each feature was centered at the median DMSO value within each plate.

### Correlation feature-weighting

Our distance-based analyses assessed the similarity of samples collectively across a range of phenotypic features. To address bias introduced by highly correlated features—i.e. highly correlated features capture similar information, which will be weighted more heavily by euclidean distance as more features pick up this information—we re-weighted l2 normalized features based on the sum of their absolute correlations. That is, we re-weighted features within each cell line as

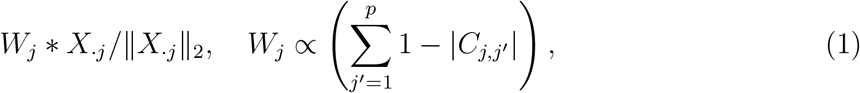

where *C*_*j,j′*_ denotes the Pearson correlation between features *j, j*^*′*^ and *X*_*·j*_ denotes feature vector *j* for all samples from a select cell line after plate position normalization.

### Replicates and multiple dose levels

In the case where the experimental data contained multiple doses of a compound, only the highest dose was taken. For each compound, we averaged position-normalized, correlation-weighted KS scores across replicates at the highest dose used (when multiple doses or replicates were present in the experimental data)

### Quantitative definitions of phenoactivity and phenosimilarity

For simplicity, the following notations are defined relative to a single cell line unless explicitly stated otherwise. Our full dataset can be viewed as *K* = 6 (cell lines) instantiations of the data described below, with the same compound library screened across each cell line. All distances described throughout this section are evaluated between phenotypic profiles as described above. Let *X ∈* R_*n×p*_ denote phenotypic profiling data for a given cell line, with rows *X*_*i*_ *∈* R_*p*_ being the phenotypic profile for sample *i*. Samples *i* = 1, …, *n* represent different compounds from a library of interest (i.e., a single replicate of a compound perturbation, which is the same as a single well on an imaging plate). Each sample has a unique compound label *m*_*i*_ *∈ {*1, …, *M }* and *n*_*m*_ denotes the number of samples labeled as MOA *m*. Let *I*_*C*_ *⊂ {*1, …, *n}* index the DMSO control samples and *I*_*m*_ *⊂ {*1, …, *n}* the samples labeled with MOA *m*.

Our goal is to select the optimal cell line(s) *S*^*∗*^ *⊂* 1, …, *K* relative to an analytical criteria of interest. Toward this end, we consider the distance between compounds in phenotypic space, with *d*_*ij*_ := *d*(*X*_*i*_, *X*_*j*_) denoting the euclidean distance between samples *i, j*. The approach described below can be used substituting another distance metric of interest with euclidean distance. Our criteria compare the distribution of distances from a query population of interest (e.g. compounds with the same MOA) to a reference population (e.g. DMSO controls). Comparing different query and reference populations allows us to address different questions. In the following sections, we show how optimal cell line selection for both phenoactivity and phenosimilarity can be framed within this context.

### Phenoactivity

Let 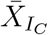 denote the median (i.e. centroid; computed feature-wise) phenotypic profile for DMSO samples. For a population of samples, indexed by *I ⊂ {*1, …, *n}*, consider their distances to the DMSO centroid

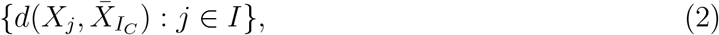

and denote the corresponding empirical cumulative distribution function (ECDF) of these distances as 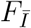. We define the phenoactivity for a MOA *m* by comparing query: *I* = *I*_*m*_ and reference: *I* = *I*_*C*_ populations as

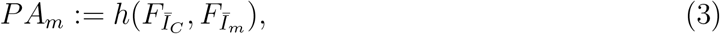

where *h* is a function measuring the deviation ECDFs. In this study, we set *h* to be a signed variant of the earth mover distance that measures the difference between ECDFs

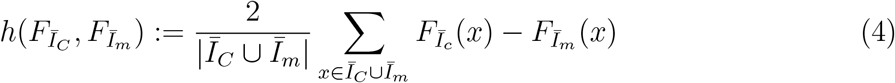

Comparing *PA*_*m*_ across cell lines allows us to evaluate which lines are most sensitive to MOA *m* (Fig 1. A-C). To evaluate phenoactivity of MOA *m* across sets of cell lines *S ⊆ {*1, …, *K}*, we take the maximum phenoactivity score over all cell lines in *S*, effectively asking whether any of these cell lines differentiates MOA *m* from DMSO control. We denote phenoactivity for a set of cell lines as

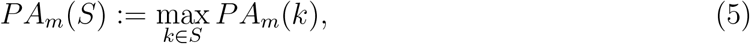

where *PA*_*m*_(*k*) is the phenoactivity score for MOA *m* in cell line *k*.
We cast cell line selection as an optimization problem

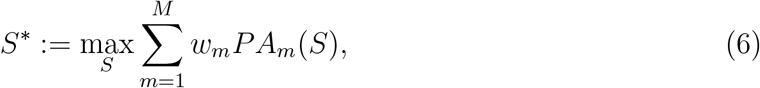

where *w*_*m*_ *∈* [0, 1] denotes a user-provided weight associated with MOA *m*. In other words, our optimization criteria is simply a weighted average of phenoactivity scores across different MOAs. Weights allow us to prioritize compound classes of interest. For instance, setting weights equal will select a “generalist” cell line that performs well across all MOAs. In contrast, setting weights higher for a particular MOAs will select cell lines that are specialists within those classes.

**Figure 1:**
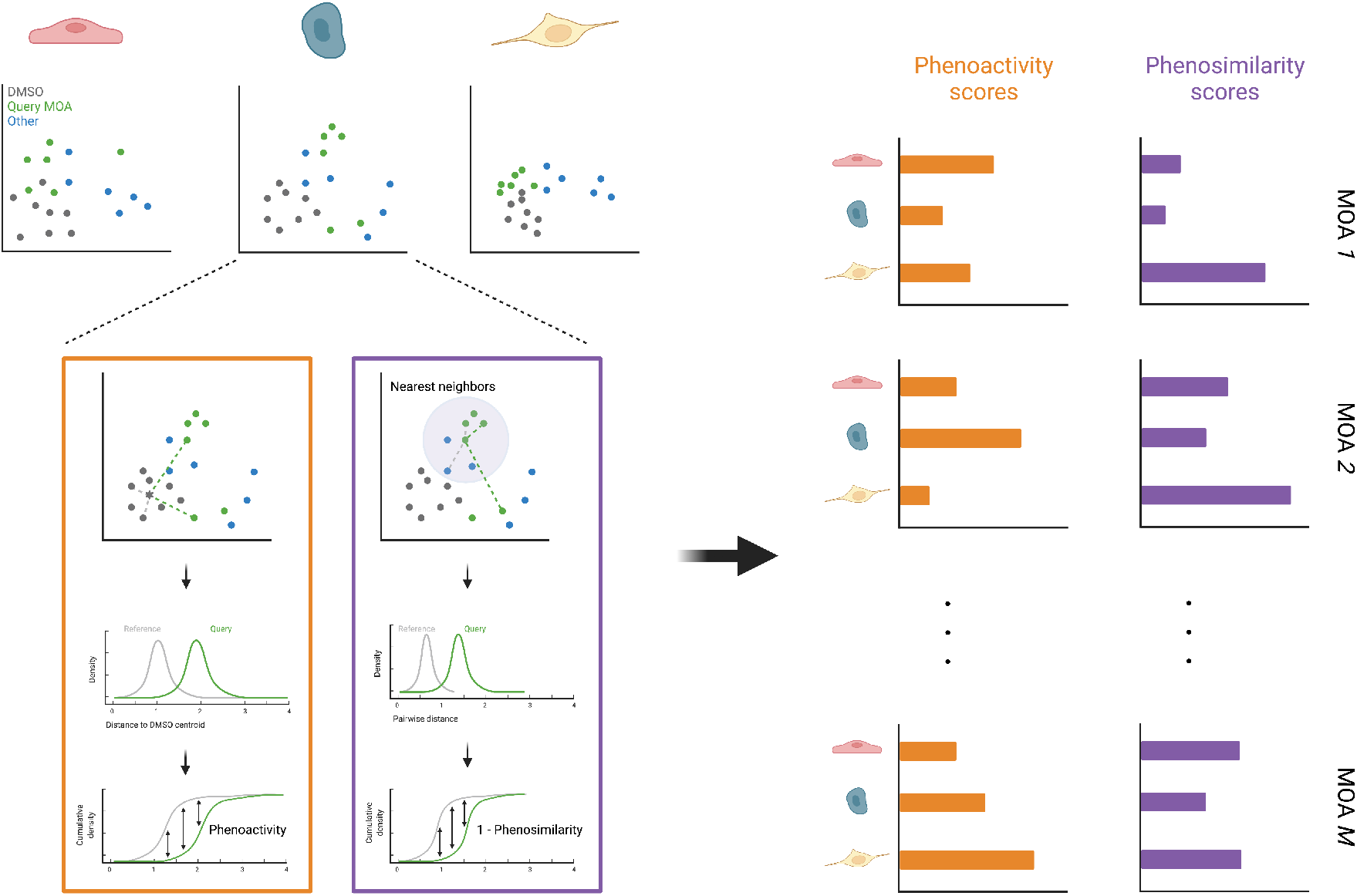
Overview of analysis. Comparing distance distributions between groups of phenotypic profiles is used to evaluate phenoactivity (orange) and phenosimilarity (purple). Phenoactivity and/or phenoactivity scores are compared across cell lines and MOAs to select optimal cell line(s).

**Figure 2:**
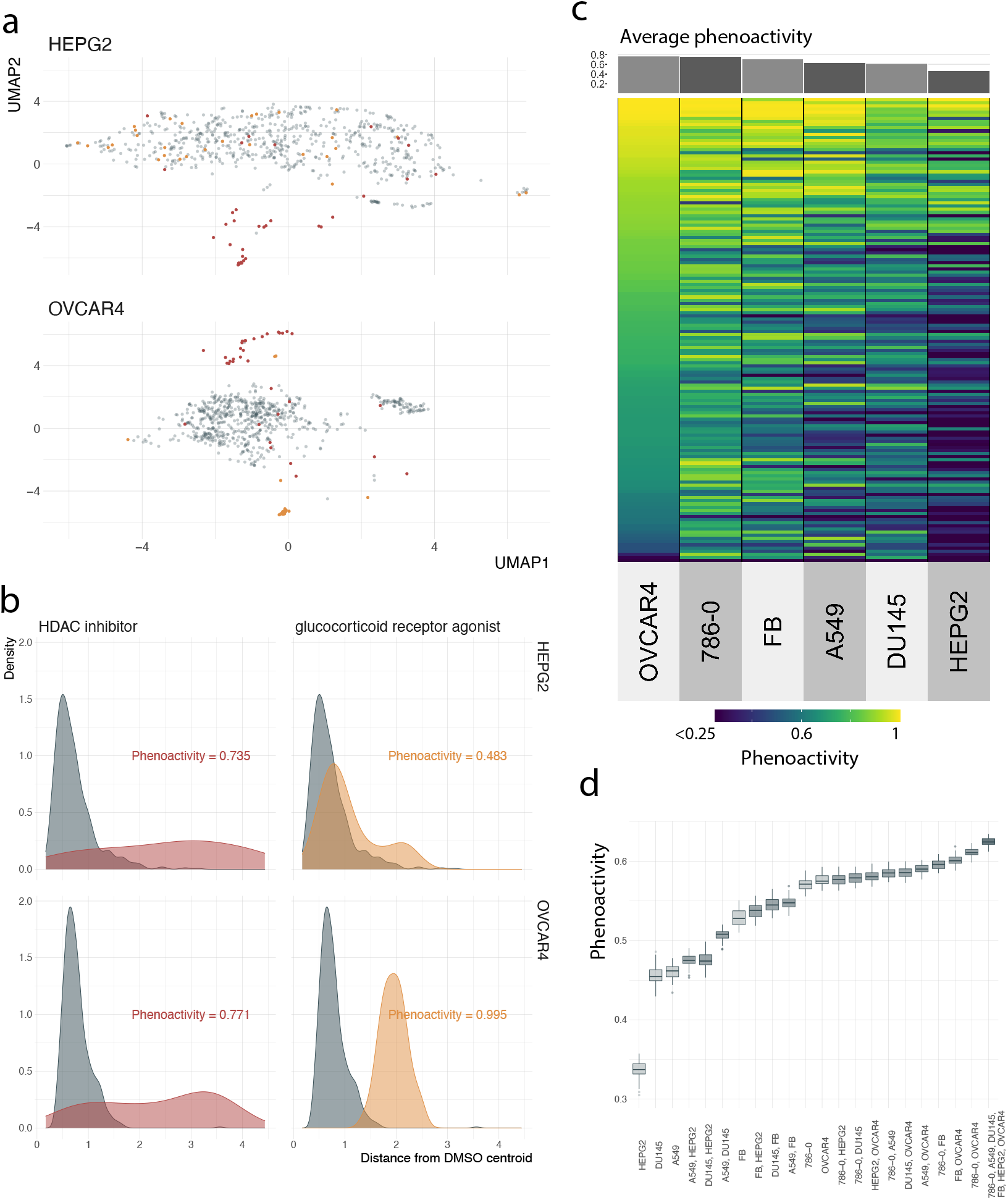
Optimizing cell line selection for phenoactivity. **(a)** UMAP projection of phenotypic profiles for query MOA samples (HEPG2 top, OVCAR4 bottom) and DMSO. **(b)** Distribution of distances to the DMSO point cloud centroid for DMSO samples and query MOAs by cell line (HEPG2 top, OVCAR4 bottom). Query distributions that are further from DMSO reference result in higher phenoactivity scores. **(a-b)** colors: compound MOA (red = HDAC inhibitor, yellow = gluccocorticoid receptor agonist). **(c)** Phenoactivity scores by cell line (column), MOA (row). MOAs are filtered to those with at least 5 distinct compounds. **(d)** Distribution of phenoactivity scores by cell line set, evaluated over 50 random subsamples of the library (2 / 3 of compounds subsampled). Color: number of cell lines from 1 (lightest) to 6 (darkest).

**Figure 3:**
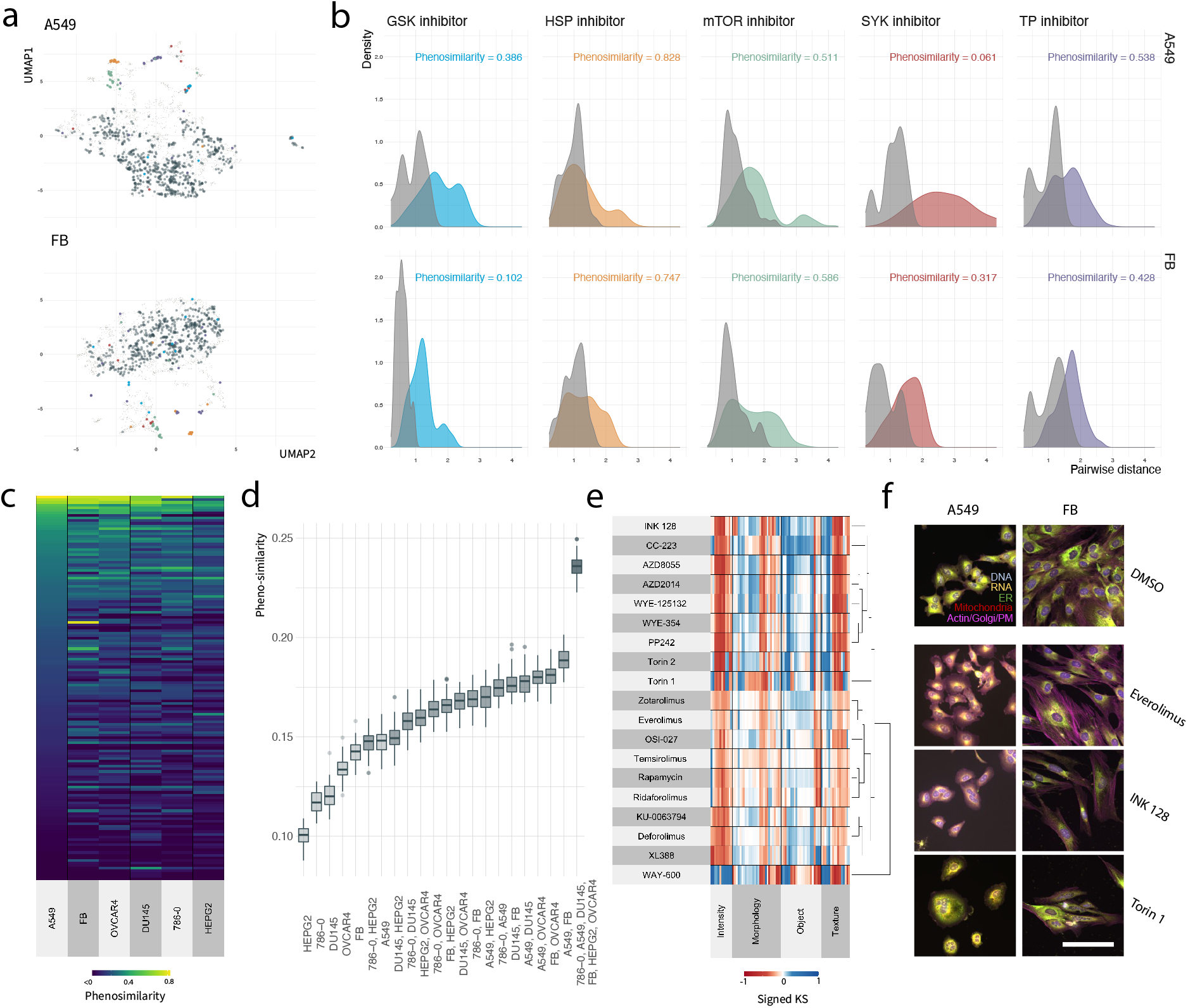
Optimizing cell line selection for phenosimilarity. **(a)** UMAP projection of phenotypic profiles by cell line (A549 top, FB bottom). **(b)** Distribution of pairwise distances between query MOA compounds (colored distribution) and their nearest neighbors (grey distribution) by cell line (A549 top, FB bottom). **(a-b)** Color: MOA (blue = GSK inhibitor, yellow = HSP inhibitor, green = mTOR inhibitor, red = SYK inhibitor, purple = TP inhibitor, dark grey = DMSO; light brown, small points = other MOA; light grey distribution corresponds to nearest neighbors of a given MOA class). **(c)** Phenosimilarity scores by cell line (column), MOA (row). MOAs are filtered to include those with at least 5 distinct compounds. **(d)** Distribution of phenoactivity scores by cell line set, evaluated over 50 random subsamples of the library (2 / 3 of compounds subsampled). Color: number of cell lines from 1 (lightest) to 6 (darkest). **(e)** Clustered phenotypic profiles for mTOR compounds evaluated in A549. **(f)** Representative images of compound treated cells (A549 or FB) after 48 hours exposure to to DMSO vehicle control 0.1% (top) and select mTOR inhibitors (bottom 3). Scale bar represents 100um.

### Phenosimilarity

We assess phenosimilarity in a query MOA *m* by evaluating whether samples are close to one another in the MOA relative to neighboring samples in phenotypic space. These neighboring samples may have the same MOA as samples *i ∈ I*_*m*_, implying that MOA *m* is tightly clustered in phenotypic space, or include different MOAs, implying that MOA *m* is intermingled with other MOAs in phenotypic space.

Consider the the pairwise distances between compounds with MOA *m {d*_*ij*_ : *i, j ∈ I*_*m*_*}* and let *F*_*Im*_ denote the corresponding ECDF. As a baseline for this distance distribution, we use the distances between samples in *I*_*m*_ and their nearest neighbors as *{d*_*ij*_ : *i ∈ I*_*m*_, *j ∈ N*_*m*_(*i*)*}*, with corresponding ECDF *F*_*Nm*_, where *N*_*m*_(*i*) are the *n*_*m*_ nearest neighbors of sample *i*. We define the phenosimilarity for MOA *m* as

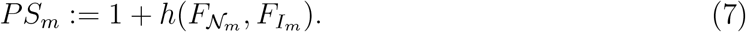

We note that *F*_*Nm*_ provides a lower bound on *F*_*Im*_ in the sense that *h*(*F*_*Nm*_, *F*_*Im*_) *≤* 0, with equality only when all nearest neighbors belong to MOA *m*. Thus (7) returns a values between 0 and 1 when all nearest neighbors are from MOA *m*; values *<* 1 represent varying degrees of overlap with other MOAs.

For a set of cell lines *S*, we define MOA similarities as the maximum across all cell lines *k ∈ S*, denoted as *PS*_*m*_(*S*). This asks whether any cell line *k ∈ S* groups MOA *m* compounds. As in the case of phenoactivity, we cast cell line selection for phenosimilarity as an optimization problem

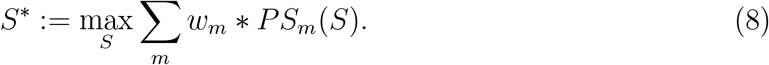

## Supporting Information

**Figure S1:**
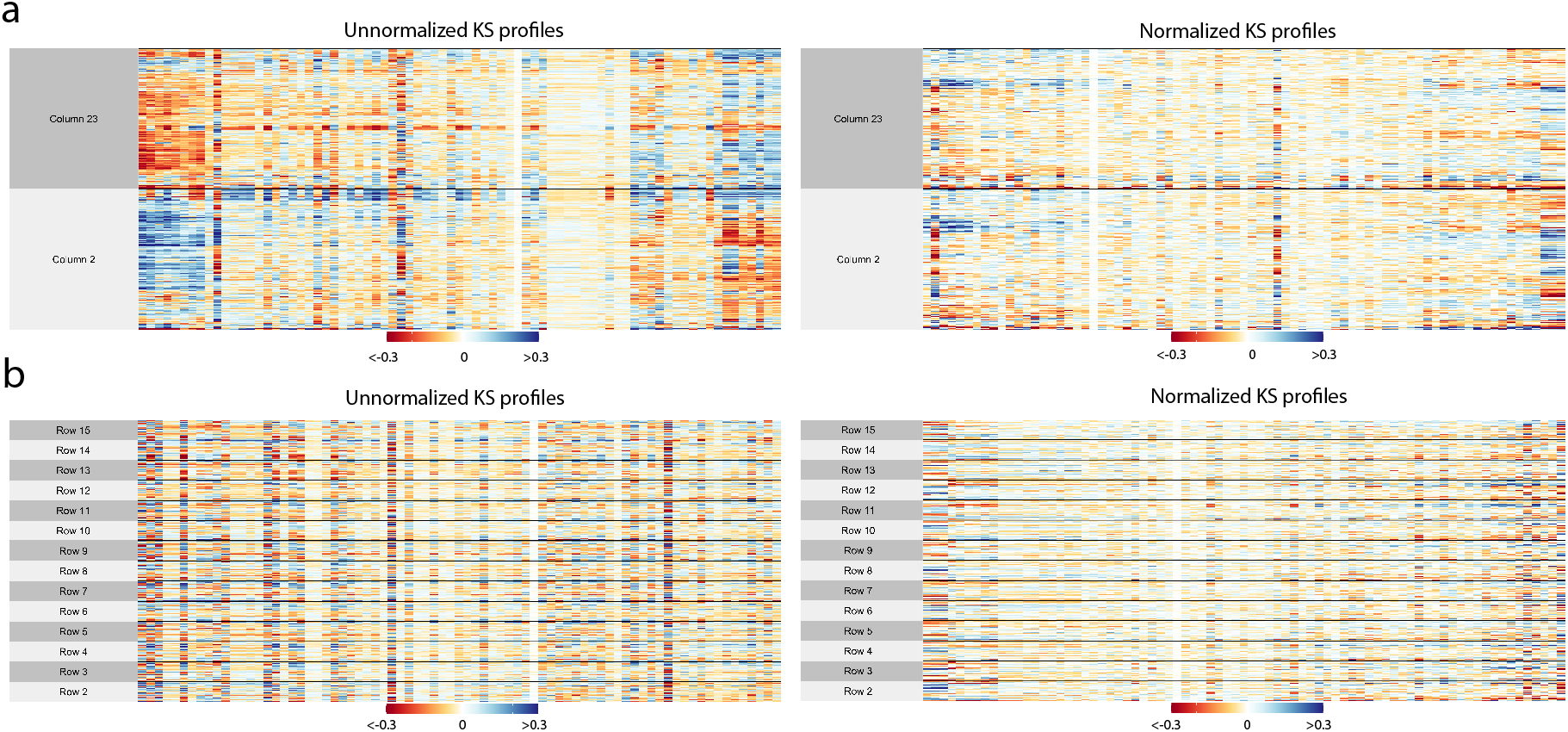
KS profiles of each DMSO sample before (left) and after (right) plate position normalization. Rows: well replicates, columns: image-derived features. **(a)** samples are grouped by column on imaging plate. **(b)** samples are grouped by row on imagining plate.

**Figure S2:**
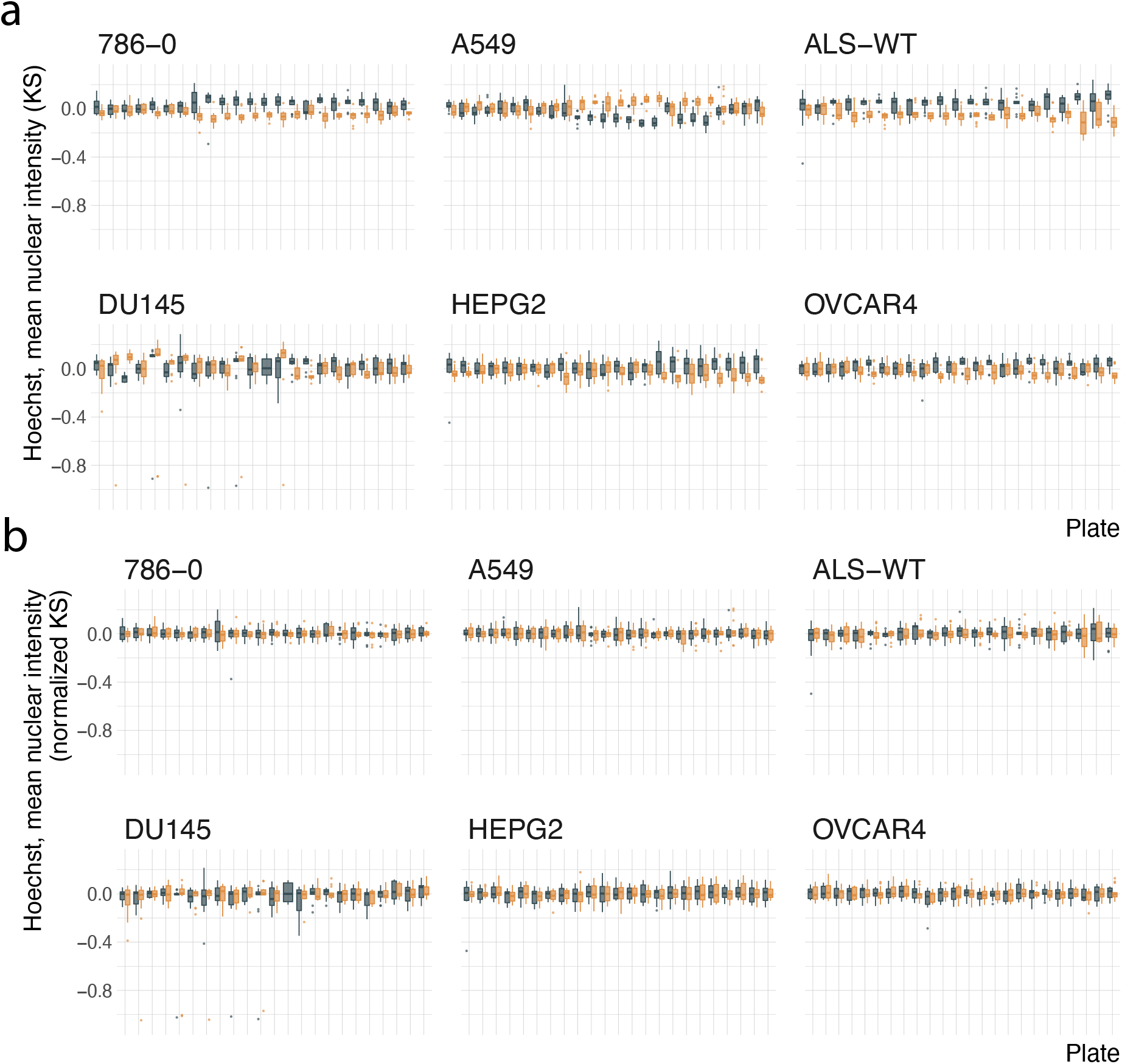
Distribution of hoechst nuclear intensity in DMSO samples before **(a)** and after **(b)** plate position normalization by plate. Color: column on imaging plate (grey = 23, yellow = 2).

**Figure S3:**
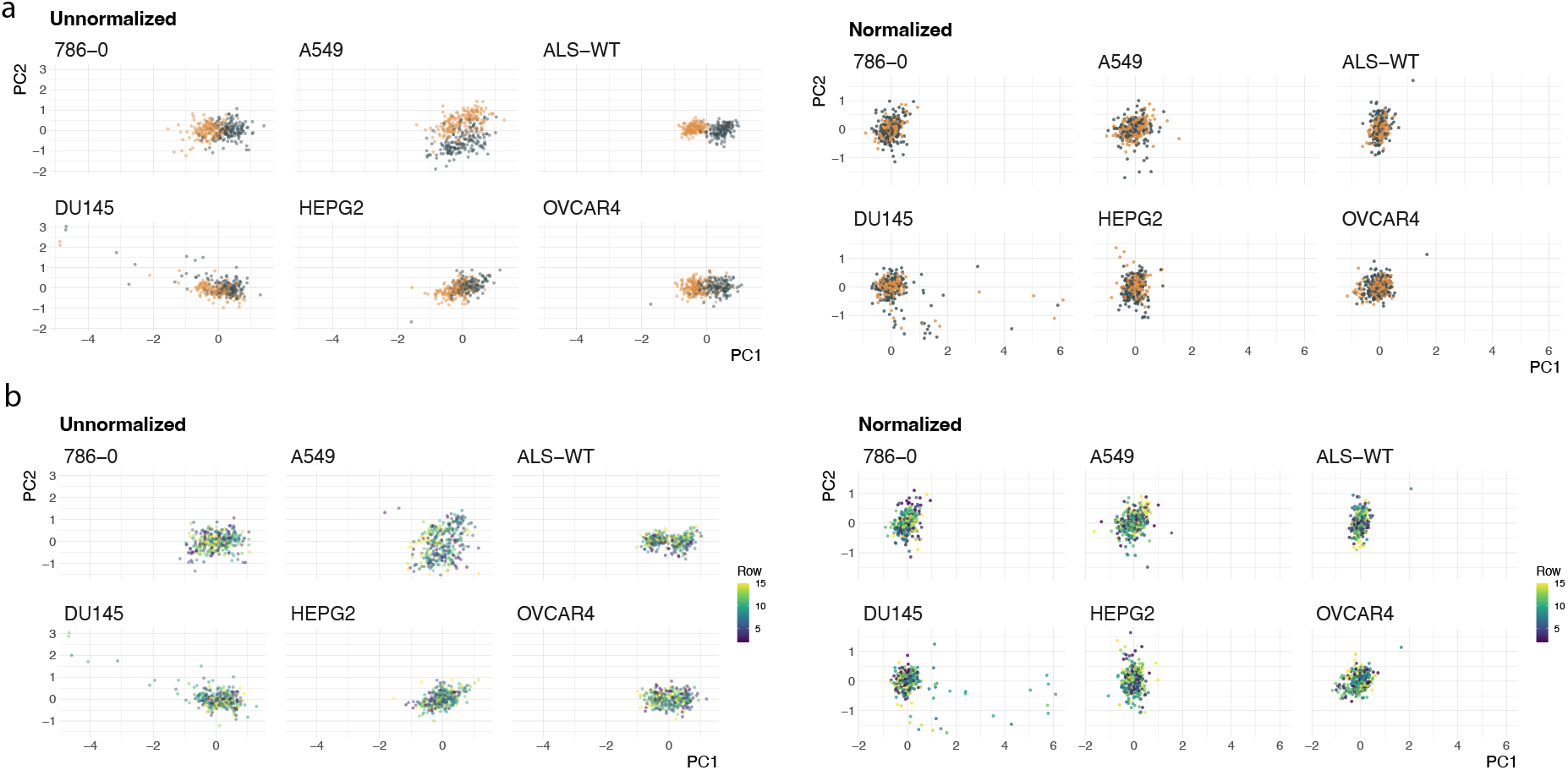
PCA projections of DMSO samples before (left) and after (right) plate position normalization. **(a)** Color: column on the imaging plate (grey = 23, yellow = 2). **(b)** Color: row on the imaging plate.

**Figure S4:**
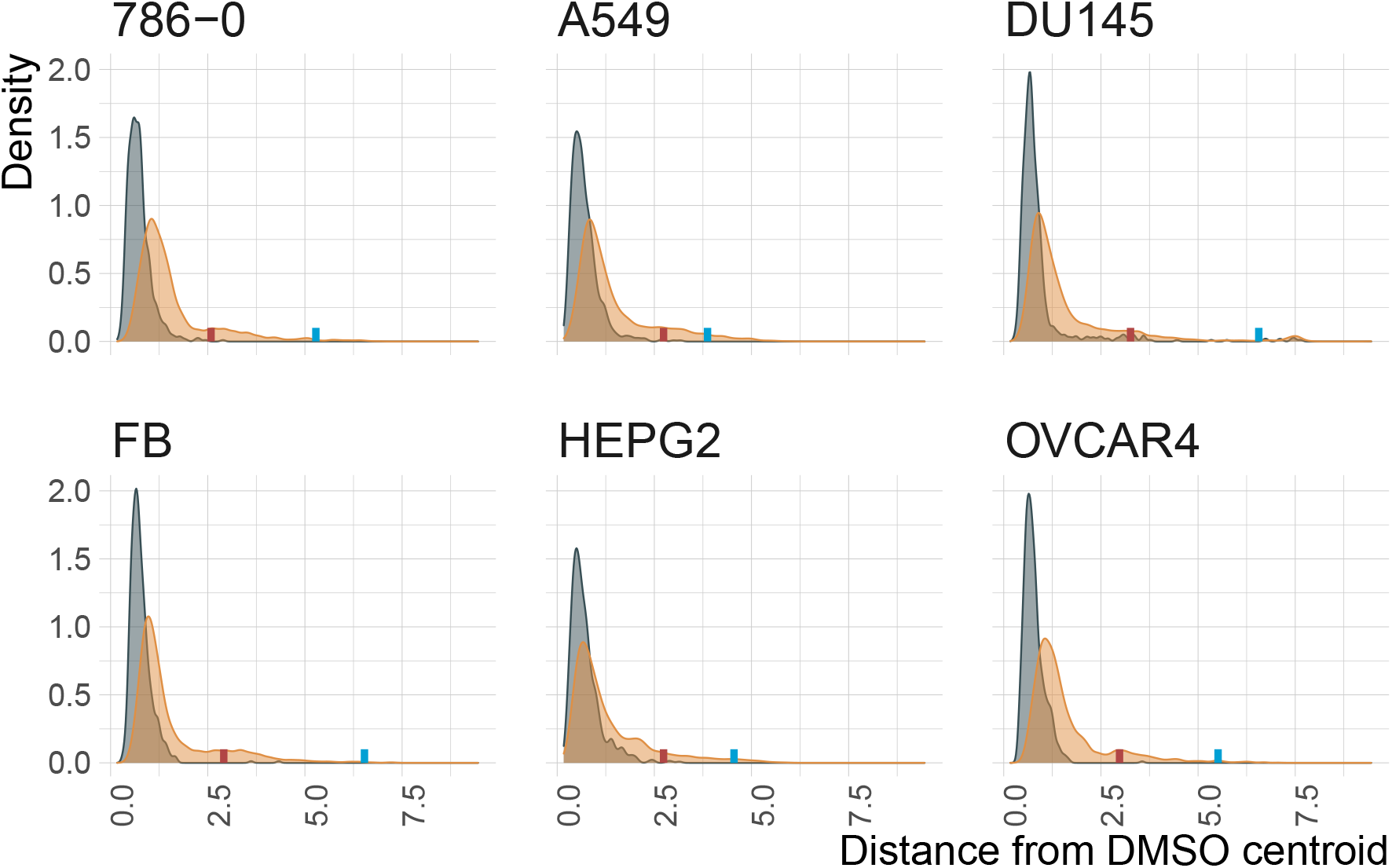
Distance to DMSO point cloud centroid by cell line. Lines indicate the distance to DMSO centroid for positive control compounds. Colors: grey = DMSO samples, yellow = query compounds; red line = Gemcitabine, blue line = Bortezomib.

**Figure S5:**
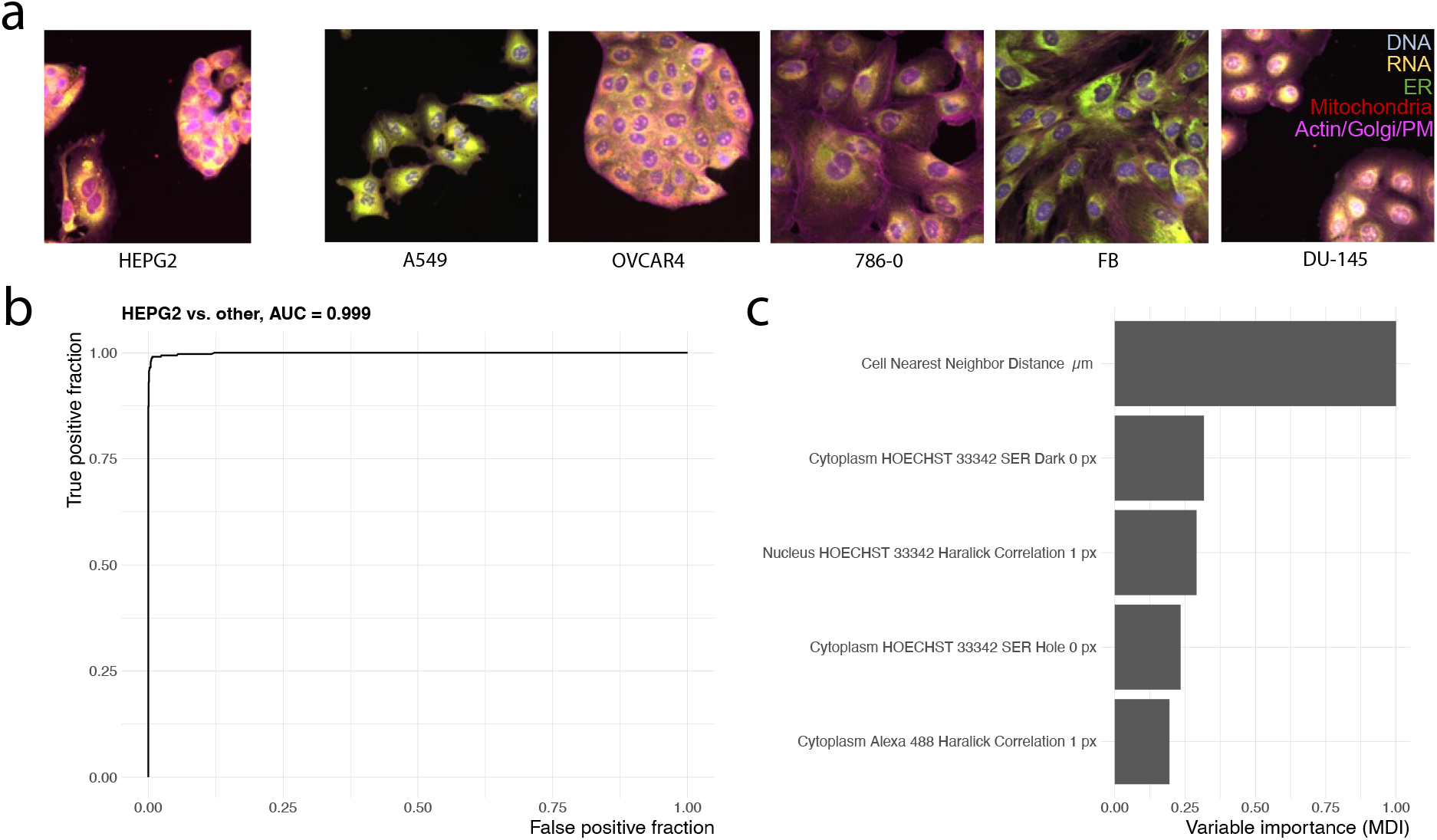
HEPG2 versus other cell lines. **(a)** Representative images of cells after 48 hours exposure to DMSO vehicle control 0.1%. Scale bar represents 100um. **(b)** ROC curve of iterative random forest model trained to classify HEPG2 versus other cell lines. **(c)** MDI feature importance of top 5 features used in HEPG2 versus other cell line classifier.

**Figure S6:**
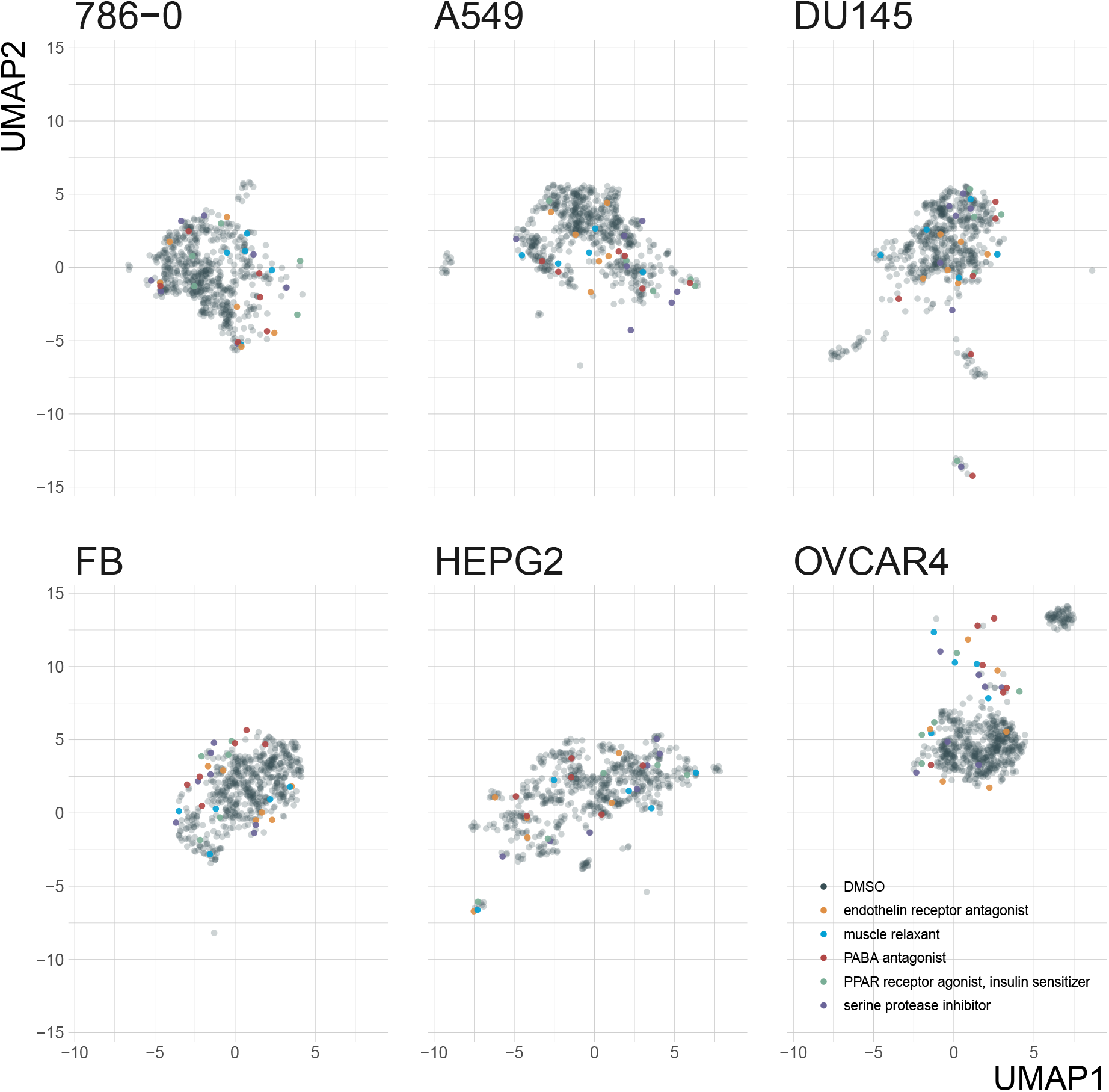
UMAP projections of MOAs with low phenosimilarity across all cell lines and DMSO controls. MOAs selected based on lowest rank of maximum phenosimilarity across all cell lines

**Figure S7:**
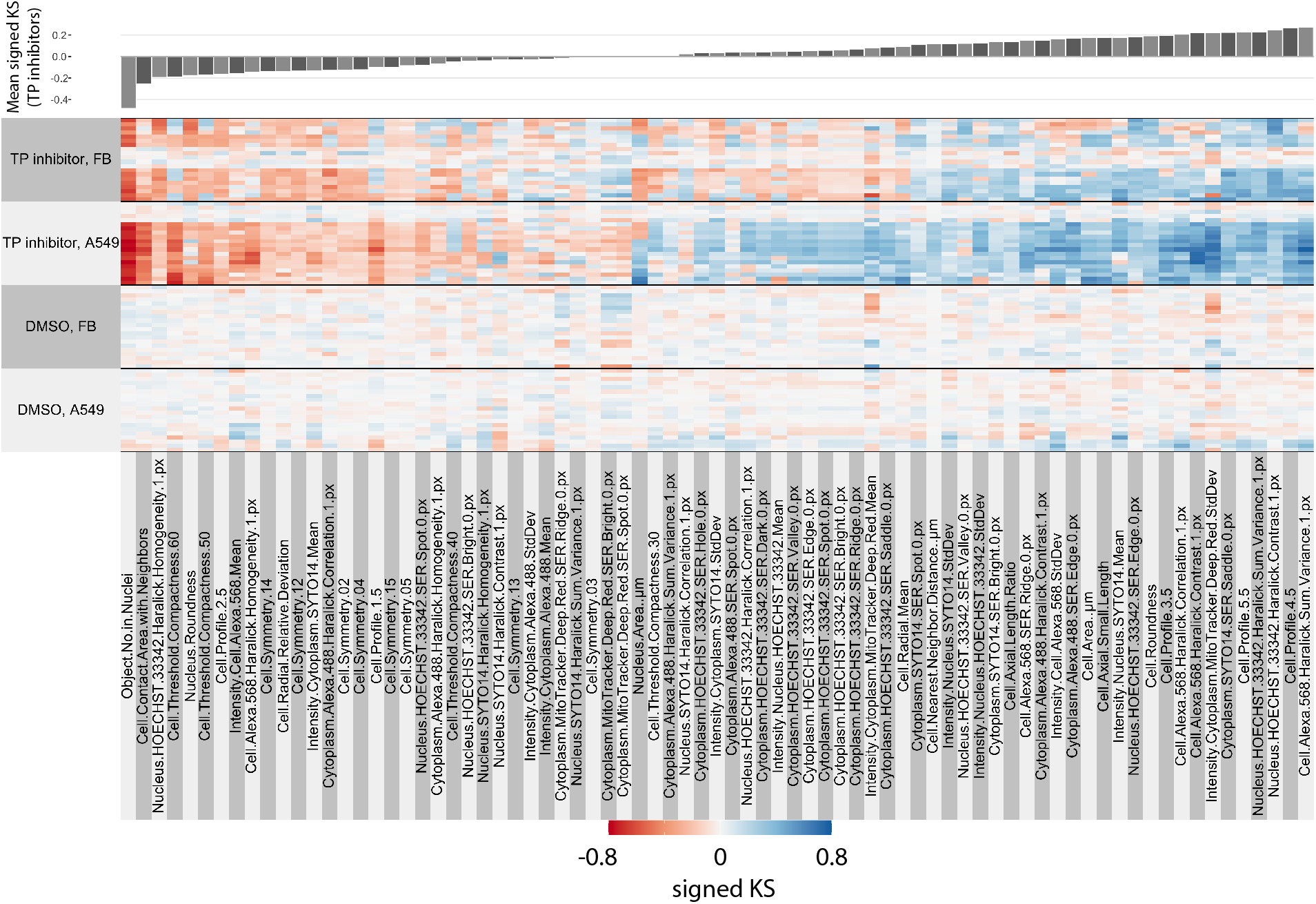
Phenotypic profiles for TP inhibitors and DMSO controls in A549 and FB cell lines. Features ordered based on average (across compounds and cell lines) signed KS value in TP inhibitors.

